# Distinct eLPB^ChAT^ projections for methamphetamine anxiety and relapse

**DOI:** 10.1101/2023.10.23.563030

**Authors:** Wenwen Chen, Hao Guo, Ning Zhou, Xing Xu, Yuning Mai, Teng He, Jun Wen, Feifei Ge, Shan Qin, Chengyong Liu, Wenzhong Wu, Hee Young Kim, Yu Fan, Xiaowei Guan

**Author notes:** Correspondence to: Xiaowei Guan, M.D., Ph.D., Yu Fan, Ph.D. Chen W, Guo H and Zhou N contributed equally to this work.

## Abstract

Choline acetyltransferase-positive neurons in the external lateral parabrachial nucleus (eLPB^ChAT^) send projections to PKCδ-positive (PKCδ^+^) neurons in lateral portion of central nucleus of amygdala (lCeA^PKCδ^) and oval portion of bed nucleus of the stria terminalis (ovBNST^PKCδ^), forming eLPB^ChAT^–lCeA^PKCδ^ and eLPB^ChAT^–ovBNST^PKCδ^ pathways. At least in part, the eLPB^ChAT^ neurons positively innervate lCeA^PKCδ^ and ovBNST^PKCδ^ through regulating synaptic elements of presynaptic acetylcholine (Ach) release and postsynaptic nicotinic acetylcholine receptors (nAChRs). Methamphetamine (METH) withdrawal anxiety and METH-primed reinstatement of conditioned place preference (CPP) recruit eLPB^ChAT^–lCeA^PKCδ^ pathway and eLPB^ChAT^–ovBNST^PKCδ^ pathway in male METH-exposed mice, respectively.

---

Methamphetamine (METH) is a highly addictive and abused psychostimulant worldwide. Among individuals with METH use disorders (MUD), 34.3% develop withdrawal anxiety symptoms ^1, 2^, and up to 90% are estimated to drive relapse ^3^, which present big challenge for clinical practice. Previously, we reported that the choline acetyltransferase-positive (ChAT^+^) neurons in the external lateral parabrachial nucleus (eLPB^ChAT^) are involved in METH-primed reinstatement of METH conditioned place preference (CPP) in mice ^4^. In the present study, we further dissected the projections of eLPB^ChAT^ neurons, and explored their roles in withdrawal anxiety and reinstatement of CPP in METH-exposed male mice.

Consistent with our previous study ^4^, we found that ChAT^+^ neurons are widely distributed in eLPB^ChAT^ along the anterior-posterior brain axis in naïve mice (Extended Data Fig. 1A-B). Our Previous findings showed that eLPB^ChAT^ neurons send direct fibers to GABAergic neurons in central nucleus of amygdala (CeA^GABA^), forming eLPB^ChAT^–CeA^GABA^ pathway. With the mapping results of fMOST ^4^, in addition to CeA, the bed nucleus of the stria terminalis (BNST) seemed to be another concentrated projecting nucleus by eLPB^ChAT^ neurons. Here, with method of anterograde (Fig. 1A) and retrograde (Extended Data Fig. 1C-D) virus tracing, we found that eLPB^ChAT^ neurons (Fig.1B) mainly projected to lateral portion of CeA (lCeA) and oval portion of BNST (ovBNST). Further, about 75% and 52% of eLPB^ChAT^ terminals were distributed around PKCб-positive (PKCδ^+^) neurons in lCeA and ovBNST, respectively, forming eLPB^ChAT^–lCeA^PKCδ^ pathway and eLPB^ChAT^–ovBNST^PKCδ^ pathway (Fig.1C).

**Fig. 1.**
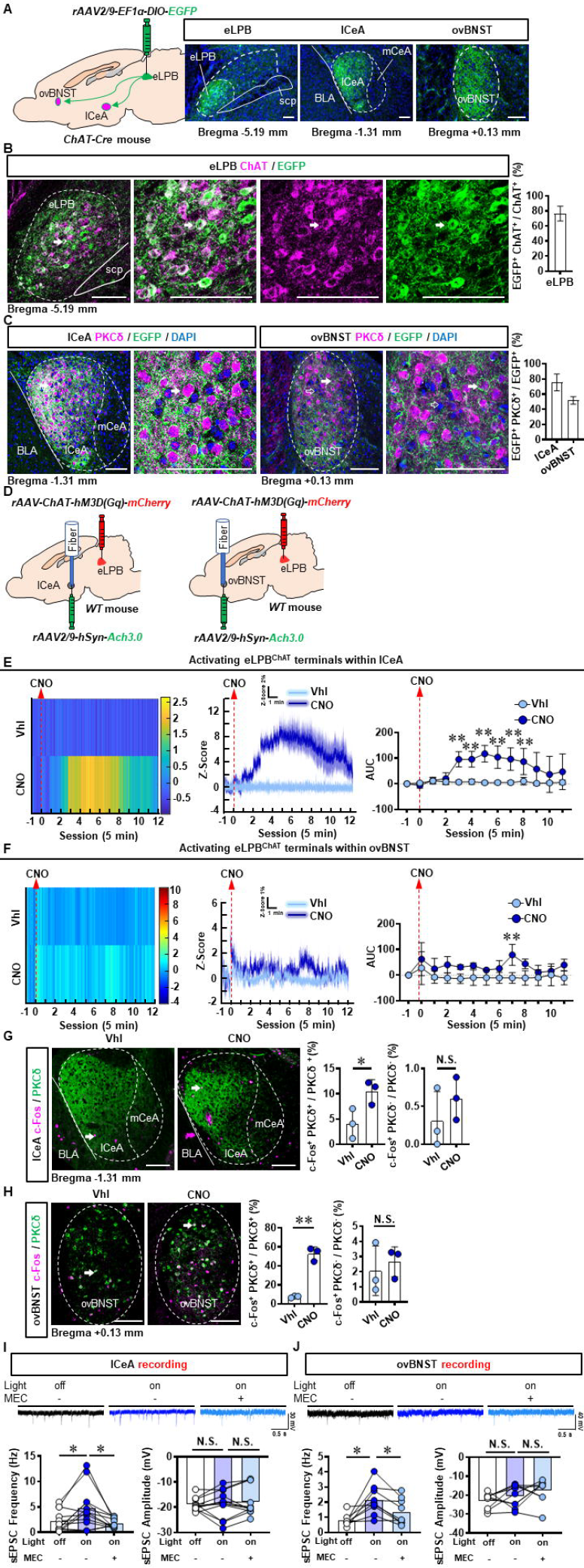
Anatomical structure and functional innervation of the eLPB^ChAT^– lCeA^PKCδ^ and eLPB^ChAT^–ovBNST^PKCδ^ pathways in naïve male mice. **A**, Schematic diagram and representative images of the *rAAV2/9-DIO-EGFP* injection in the eLPB, lCeA and ovBNST in ChAT-Cre mouse. Scale bar, 100 μm. **B**, Representative images and the percentage of EGFP-labelled neurons in eLPB ChAT-positvie populations. **C**, Representative images and the percentage of EGFP-labelled terminals that surrounded PKCδ-positive (PKCδ^+^) neurons in the lCeA and ovBNST. **D**, Schematic diagram of *rAAV-ChAT-hM3D(Gq)-mCherry* injection into eLPB and fiber implantation in lCeA or ovBNST in wild-tye (WT) mouse. **E**, Heatmap (left), quantification (middle) and AUC (right) of Z-Score of Ach3.0 fluorescence in lCeA. Two-way ANOVA with Sidak’s multiple comparisons test. n = 5 mice per group. F _(12, 104)_ = 4.730, *p* < 0.0001; t = 4.545, \**p* (15 min) = 0.0002; t = 4.482, \**p* (20 min) = 0.0002; t = 5.780, \*\**p* (25 min) < 0.0001; t = 5.069, \**p* (30 min) < 0.0001; t = 4.675, \**p* (35 min) = 0.0001; t = 3.900, \**p* (40 min) = 0.0022. **F**, Heatmap (left), quantification (middle) and AUC (right) of Z-Score of Ach3.0 fluorescence in ovBNST. Two-way ANOVA with Sidak’s multiple comparisons test. n = 3 mice per group. F _(12, 52)_ = 0.7841, *p* = 0.6640; t = 3.613, \*\**p* (35 min) = 0.0088. **G**, The percentage of c-Fos-positive (c-Fos^+^) neurons in lCeA PKCδ^+^ and PKCδ-negative (PKCδ^-^) neurons. Two-tailed unpaired t test. n = 3 mice per group. c-Fos^+^ PKCδ^+^, t _(4)_ = 2.925, *p* = 0.043; c-Fos^+^ PKCδ^-^, t _(4)_ = 1.076, *p* = 0.3426. **H**, The percentage of c-Fos^+^ neurons in ovBNST PKCδ^+^ and PKCδ^-^ neurons. Two-tailed unpaired t test. n = 3 mice per group. c-Fos^+^ PKCδ^+^, t _(4)_ = 10.89, *p* = 0.0004; c-Fos^+^ PKCδ^-^, t _(4)_ = 0.5378, *p* = 0.6192. **I**, Representative traces and statistics data of lCeA neurons sEPSC. n = 12 cells from 6 mice. The one-way analysis of variance (ANOVA) with Tukey post-test. Frequency, F _(1.440, 15.84)_ = 7.384, *p* = 0.0093; \**p* = 0.0316 off vs on; \**p* = 0.0365 on vs on-MEC; Amplitude, F _(1.825, 20.07)_ = 0.4156, *p* = 0.6473; *p* = 0.9690 off vs on; *p* = 0.7725 on vs on-MEC. **J**, Representative traces and statistics data of ovBNST neurons sEPSC. n = 12 cells from 6 mice. The one-way analysis of variance (ANOVA) with Tukey post-test. Frequency, F _(1.486, 11.89)_ = 9.790, *p* = 0.0049; * *p* = 0.0156 off vs on; \**p* = 0.0115 on vs on-MEC; Amplitude, F _(1.859, 14.87)_ = 2.762, *p* = 0.0983; *p* = 0.5846 off vs on; *p* = 0.4936 on vs on-MEC. Vhl, vehicle; CNO, clozapine-N-oxide. *, *p* < 0.05, ** *p* < 0.01 vs Vhl. Scale bar, 100 μm.

Since cholinergic neurons could play the role of interneuron in local regulation ^5^ or innervating other nuclei through long projections ^6, 7^ in the brain, we explored the physiological innervation of eLPB^ChAT^ neurons on PKCδ^+^ neurons in lCeA and ovBNST. First, to observe the real-time acetylcholine (Ach) signals from eLPB^ChAT^ terminals within lCeA and ovBNST, designer receptors exclusively activated by designer drugs (DREADDs) method and Ach fiber photometry were used in male naïve mice (Fig. 1D, Extended Data Fig. 2A-B). As shown in Extended Data Fig. 2C, systemic clozapine-N-oxide (CNO) administration efficiently activate eLPB^ChAT^ neurons. In parallel, the Ach release was significantly increased in lCeA at 15 min to 40 min (Fig. 1E) and in ovBNST at 35 min after CNO injection (Fig. 1F) in free-moving male mice, when compared with vehicle controls. In another set of same mice models, we collected the brain tissue at 35 min after CNO injection, and found that both the lCeA^PKCδ^ neurons (Fig. 1G) and ovBNST^PKCδ^ neurons (Fig. 1H) were evoked by activating eLPB^ChAT^ neurons, as indicated by increased c-Fos^+^ PKCδ^+^ neurons. Next, to characterize the potential postsynaptic elements for the eLPB^ChAT^–lCeA/ovBNST pathway, we combined optogenetics activation strategy and patch-clamp recording in acutely prepared slices (Extended Data Fig. 2D-E). The blue light (473 nm) was used to activate the eLPB^ChAT^ terminals within lCeA or ovBNST. As shown in Fig. 2I-J, the frequency of spontaneous excitatory postsynaptic currents (sEPSC) in the lCeA and ovBNST neurons was increased when activating LPB^ChAT^ terminals. When inhibiting nicotinic acetylcholine receptors (nAChRs) with mecamylamine incubation during optogeneticly activating eLPB^ChAT^ terminals, the triggered sEPSC frequency were blocked in slices of lCeA and ovBNST. In parallel, the amplitude of sEPSC has no significant difference in the lCeA and ovBNST neurons in the whole processes. Together, these findings suggest that eLPB^ChAT^ neurons positively excite lCeA^PKCδ^ and ovBNST^PKCδ^ neurons, at least in part being through synaptic elements of presynaptic Ach release and postsynaptic nAChRs.

**Fig. 2.**
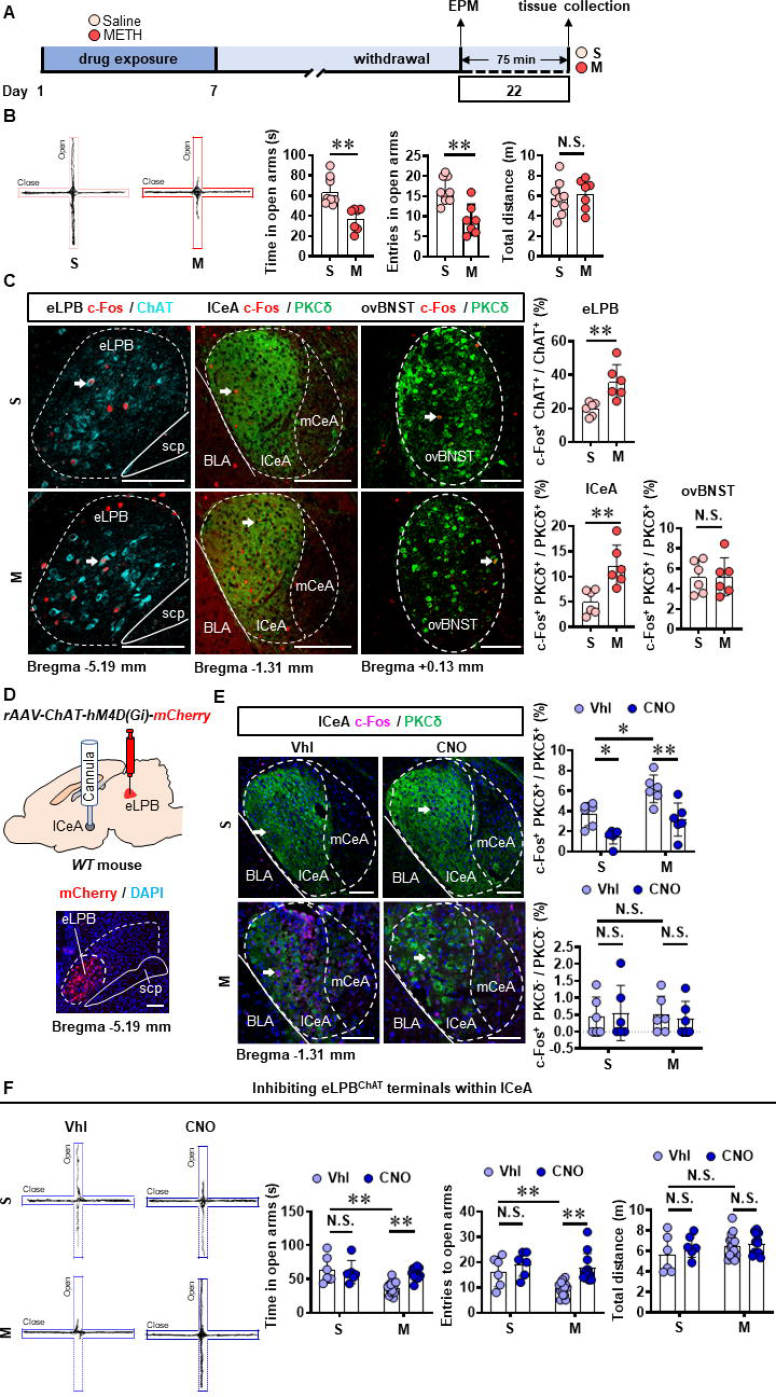
Inhibiting eLPB^ChAT^–lCeA^PKCδ^ pathway alleviates the anxiety-like behaviors in METH-withdrawn male mice. **A**, Experimental design and timeline. **B**, EPM test. Two-tailed unpaired t test. S group, n = 9 mice; M group, n = 7 mice. Time in open arms, t _(14)_ = 4.060, *p* = 0.0012. Entries into open arms, t _(14)_ = 4.194, *p* = 0.0009. Total distance, t _(14)_ = 0.4946, *p* = 0.6285. **C**, Representative images and the percentage of c-Fos^+^ neurons in eLPB^ChAT^, lCeA^PKCδ^ and ovBNST^PKCδ^ neurons. Two-tailed unpaired t test. n = 6 mice per group. eLPB, t _(10)_ = 3.539, *p* = 0.0054; lCeA, t _(10)_ = 3.636, *p* = 0.0046; ovBNST, t _(10)_ = 0.02581, *p* = 0.9799. **D**, Schematics of *rAAV-ChAT-hM4D(Gi)-mCherry* injection in eLPB and cannula implantation in lCeA of WT mouse. **E**, Representative images and the percentage of c-Fos^+^ neurons in and lCeA PKCδ^+^ and PKCδ^-^ neurons. Two-way ANOVA with Sidak’s multiple comparisons test. Upper, n = 6 mice per group. F _(1, 20)_ = 0.6286, *p* = 0.4372; S-CNO group, t = 3.024, *p* = 0.0395 vs S-Vhl group; M-CNO group, t = 4.145, *p* = 0.0030 vs M-Vhl group; M-Vhl group, t = 3.405, *p* = 0.0167 vs S-Vhl group. Lower, n = 6 mice per group. F _(1, 20)_ = 0.2019, *p* = 0.6580; S-CNO group, t = 0.2862, *p* = 0.9999 vs S-Vhl group; M-CNO group, t = 0.3492, *p* = 0.9996 vs M-Vhl group; M-Vhl group, t = 0.1561, *p* > 0.9999 vs S-Vhl group. **F**, EPM test. Two-way ANOVA with Sidak’s multiple comparisons test. S-Vhl group, n = 6 mice; S-CNO group, n = 6 mice; M-Vhl group, n = 16 mice; M-CNO group, n = 12 mice. Time in open arms, F _(1, 36)_ = 7.915, *p* = 0.0079; S-CNO group, t = 0.5496, *p* = 0.9950 vs S-Vhl group; M-CNO group, t = 4.269, *p* = 0.0008 vs M-Vhl group; M-Vhl group, t = 4.552, *p* = 0.0004 vs S-Vhl group. Entries into open arms, F _(1, 36)_ = 2.688, *p* = 0.1098; S-CNO group, t = 1.132, *p* = 0.8423 vs S-Vhl group; M-CNO group, t = 4.684, *p* = 0.0002 vs M-Vhl group; M-Vhl group, t = 3.054, *p* = 0.0251 vs S-Vhl group. Total distance, F _(1, 36)_ = 0.5195, *p* = 0.4757; S-CNO group, t = 1.138, *p* = 0.8395 vs S-Vhl group; M-CNO group, t = 0.4135, *p* = 0.9990 vs M-Vhl group; M-Vhl group, t = 1.446, *p* = 0.6409 vs S-Vhl group. Vhl, vehicle; CNO, clozapine-N-oxide; S, saline; M, methamphetamine; N.S., *p* > 0.05, *, *p* < 0.05, **, *p* < 0.01 vs S or Vhl or CNO. Scale bar, 100 μm.

The pathways of LPB–CeA and LPB–BNST have been implicated in anxiety ^8-11^. Here, we found that both eLPB^ChAT^ neurons and lCeA^PKCδ^ neurons but not ovBNST^PKCδ^ neurons were more triggered in METH-withdrawn male mice, accompanying by increased anxiety-like behaviors (Fig. 2A-C). Locally inhibiting eLPB^ChAT^ terminals within lCeA with viral tools (Extended Data Fig. 3A, Fig. 2D) efficiently attenuated the activities of PKCδ^+^ neurons but had no effect on that of PKCδ^-^ neurons in lCeA of METH-withdrawn and controlled mice (Fig. 2E). Importantly, the suppression of eLPB^ChAT^–lCeA^PKCδ^ pathway efficiently attenuate the anxiety-like behaviors in METH-withdrawn male mice, but had no influence on the related behaviors in controls (Fig. 2F). In addition, when activating eLPB^ChAT^ terminals within lCeA with viral tools (Extended Data Fig. 3B-C), the PKCδ^+^ but not PKCδ^-^ neurons were more activated in lCeA (Extended Data Fig. 3D). The eLPB^ChAT^–lCeA^PKCδ^ pathway activation induces more anxiety-like behaviors in METH-withdrawn mice, and has no influence on that in controlled mice (Extended Data Fig. 3E). The limitation of current study is that we could not specifically regulate CeA^PKCδ^ or BNST^PKCδ^ neurons *in vivo* since lacking PKCб-promotor-tagged viral tools. Together, these results suggest that eLPB^ChAT^–lCeA^PKCδ^ pathway be involved in coding METH withdrawal anxiety, while has no influence on normal anxiety-like behaviors in controls.

**Fig. 3.**
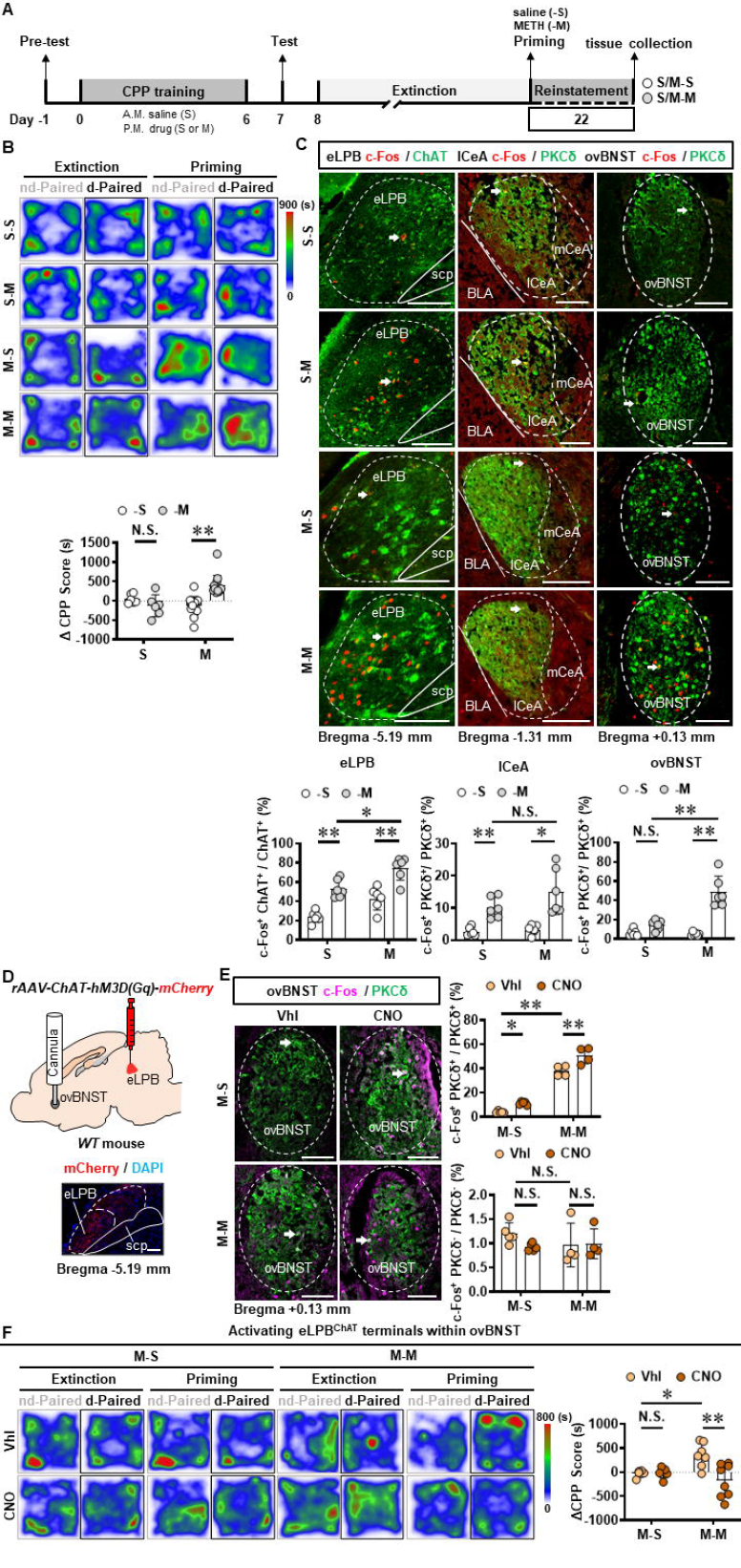
Activating eLPB^ChAT^–ovBNST^PKCδ^ pathway suppresses METH-primed reinstatement of CPP in male METH-exposed mice. **A**, Experimental design and timeline. **B**, Heatmap of spent duration by mice in CPP apparatus and ∆CPP score. Two-way ANOVA with Sidak’s multiple comparisons test. S group, n = 6 mice per group; M group, n = 14 mice per group. F _(1, 36)_ = 18.16, *p* = 0.0001; S-M group, t = 1.267, *p* = 0.7630 vs S-S group; M-M group, t = 5.846, *p* < 0.0001 vs M-S group. **C**, Representative images and the percentage of c-Fos^+^ neurons in eLPB^ChAT^, lCeA^PKCδ^ and ovBNST^PKCδ^ neurons. Two-way ANOVA with Sidak’s multiple comparisons test. n = 6 mice per group. eLPB, F _(1, 20)_ = 0.04656, *p* = 0.8313; S-M group, t = 4.967, *p* = 0.0004 vs S-S group; M-M group, t = 5.272, *p* = 0.0002 vs M-S group; M-M group, t = 3.527, *p* = 0.0126 vs S-M group. lCeA, F _(1, 20)_ = 1.737, *p* = 0.2024; S-M group, t = 3.183, *p* = 0.0277 vs S-S group; M-M group, t = 5.047, *p* = 0.0004 vs M-S group; M-M group, t = 2.149, *p* = 0.2367 vs S-M group. ovBNST, F _(1, 20)_ = 24.78, *p* < 0.0001; S-M group, t = 1.587, *p* = 0.5608 vs S-S group; M-M group, t = 8.627, *p* < 0.0001 vs M-S group; M-M group, t = 6.764, *p* < 0.0001 vs S-M group. **D**, Schematics of *rAAV-ChAT-hM3D(Gq)-mCherry* injection in eLPB and cannula implantation in ovBNST of WT mouse. **E**, Representative images and the percentage of c-Fos^+^ neurons in ovBNST lCeA PKCδ^+^ and PKCδ^-^ neurons. Two-way ANOVA with Sidak’s multiple comparisons test. Upper, M-S group, n = 5 mice per group; M-M group, n = 4 mice per group. F _(1, 14)_ = 2.226, *p* = 0.1579; M-S-CNO group, t = 3.091, *p* = 0.0469 vs M-S-Vhl group; M-M-CNO group, t = 4.766, *p* = 0.0018 vs M-M-Vhl group; M-M-Vhl group, t = 13.42, *p* < 0.0001 vs M-S-Vhl group. Lower, M-S group, n = 5 mice per group; M-M group, n = 4 mice per group. F _(1, 14)_ = 1.312, *p* = 0.2713; M-S-CNO group, t = 1.562, *p* = 0.5972 vs M-S-Vhl group; M-M-CNO group, t = 0.1395, *p* > 0.9999 vs M-M-Vhl group; M-M-Vhl group, t = 1.270, *p* = 0.7828 vs M-S-Vhl group. **F**, Heatmap of spent duration by mice in CPP apparatus and ∆CPP score (priming CPP score minus extinction CPP score). Two-way ANOVA with Sidak’s multiple comparisons test. M-S-Vhl group, n = 6 mice; M-S-CNO group, n = 6 mice; M-M-Vhl group, n = 7 mice; M-M-CNO group, n = 8 mice. F _(1, 23)_ = 8.648, *p* = 0.0073; M-S-CNO group, t = 0.1133, *p* > 0.9999 vs M-S-Vhl group; M-M-CNO group, t = 4.279, *p* = 0.0017 vs M-M-Vhl group; M-M-Vhl group, t = 2.887, *p* = 0.0489 vs M-S-Vhl group. Vhl, vehicle; CNO, clozapine-N-oxide; S-S, saline challenge-primed reinstatement test following saline CPP extinction training; S-M, METH challenge-primed reinstatement test following saline CPP extinction training; M-S, saline-primed reinstatement test following METH CPP extinction training; M-M, METH-primed reinstatement test following METH CPP extinction training; N.S., *p* > 0.05, *, *p* < 0.05, **, *p* < 0.01 vs S-S or M-S or Vhl or CNO. Scale bar, 100 μm.

Recently, emerging evidence demonstrate that PKCδ^+^ GABAergic neurons might involve in encoding relapse to drugs ^12, 13^. Here, we found that a single METH challenge (0.5 mg/kg) effectively primed the reinstatement of CPP in METH-exposed mice rather than saline-exposed mice (Fig. 3A-B, Extended Data Fig. 4A), along with increased activities of eLPB^ChAT^, lCeA^PKCδ^ and ovBNST^PKCδ^ neurons in METH-primed mice (Fig. 3C). With the phenotypes of neuronal activation in METH-primed mice in the current study, we speculate that precisely inhibiting eLPB^ChAT^–lCeA^PKCδ^ or eLPB^ChAT^–ovBNST^PKCδ^ pathway could slow down the METH-primed reinstatement of CPP in METH-exposed mice. As shown in Extended Data Fig. 5A-C, using viral tools combined with local DREADDs method, we successfully inhibiting eLPB^ChAT^ terminals in lCeA^PKCδ^ or ovBNST^PKCδ^ in METH-exposed mice. On the contrary to the expectation, neither inhibiting eLPB^ChAT^–lCeA^PKCδ^ pathway (Extended Data Fig. 5D) nor inhibiting eLPB^ChAT^–ovBNST^PKCδ^ pathway (Extended Data Fig. 5E) has influence on METH-primed reinstatement of CPP in mice. Further, there was no significances in ∆CPP score between vehicle-treated and CNO-treated groups in saline-challenged METH-exposed mice (Extended Data Fig. 5D-E), indicating that inhibiting eLPB^ChAT^ terminals in lCeA^PKCδ^ or ovBNST^PKCδ^ without administrating METH challenge can not prime the reinstatement of CPP in mice. Previously, we found that it is activating eLPB^ChAT^ neurons or CeA–projecting eLPB^ChAT^ neurons instead of suppressing them could block METH-primed reinstatement of CPP in mice ^4^. As such, we then locally activating eLPB^ChAT^ terminals in lCeA^PKCδ^ (Extended Data Fig. 6A-C) or ovBNST^PKCδ^ (Fig. 3D-E) in METH-exposed mice. We found that it was activating eLPB^ChAT^–ovBNST^PKCδ^ pathway (Fig. 3F, Extended Data Fig. 6D) rather than eLPB^ChAT^–lCeA^PKCδ^ pathway (Extended Data Fig. 6E) that block METH-primed reinstatement of METH CPP in mice. We thought that METH priming-activated eLPB^ChAT^ neurons might be a compensatory response of the brain, which possibly weaken sensitivity to drug priming through inhibiting ovBNST^PKCδ^ neurons. As shown in Fig. 3C, METH priming triggered more eLPB^ChAT^ and ovBNST^PKCδ^ neurons but not affect the lCeA^PKCδ^ neurons in METH-exposed mice than in saline-exposed mice, implying that ovBNST^PKCδ^ neurons is more involved in METH-primed reinstatement of METH CPP. Taken together, these results imply that eLPB^ChAT^–ovBNST^PKCδ^ pathway is involved in METH-primed relapse.

**Fig. 4.**
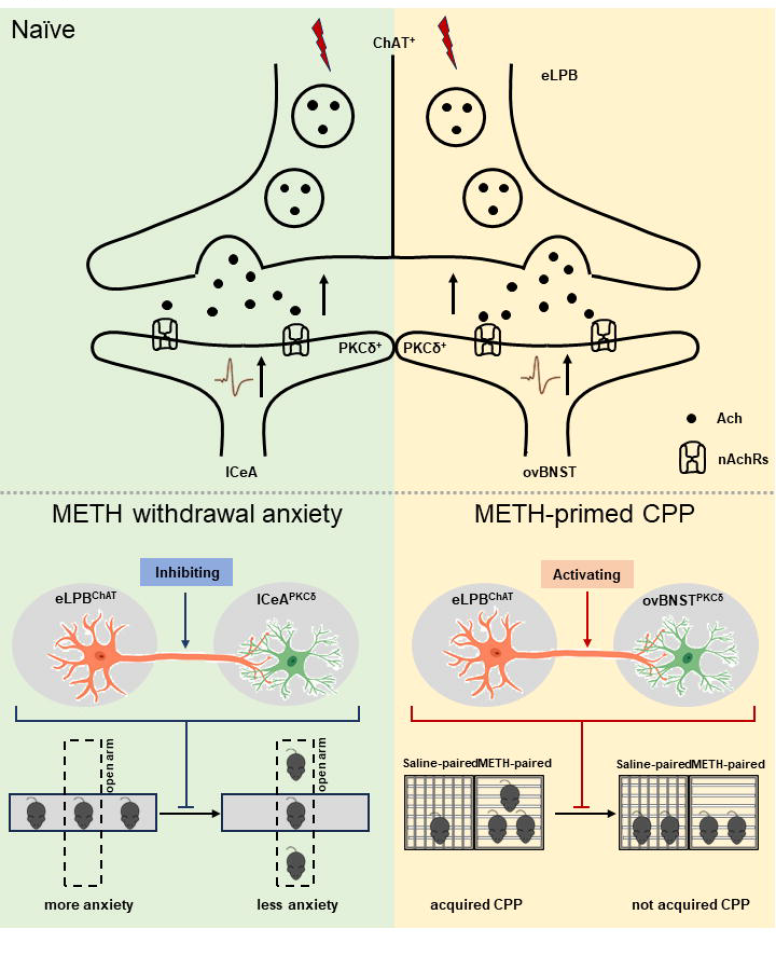
Schematic diagram of the present study. The ChAT^+^ neurons in the eLPB send projections to PKCδ^+^ neurons in lCeA and ovBNST, forming eLPB^ChAT^–lCeA^PKCδ^ and eLPB^ChAT^–ovBNST^PKCδ^ pathways. At least in part, the eLPB^ChAT^ neurons positively excite lCeA^PKCδ^ neurons and ovBNST^PKCδ^ neurons through synaptic elements of presynaptic Ach release and postsynaptic nAChRs. Chemogenetic inhibiting the eLPB^ChAT^ terminals within lCeA alleviates the anxiety-like behaviors in METH-withdrawn mice, and chemogenetic activating eLPB^ChAT^ terminals within ovBNST blocks METH-primed reinstatement of CPP in METH-exposed mice. These results indicate that METH withdrawal anxiety and METH-primed relapse recruit distinct eLPB^ChAT^ projections, as identified by eLPB^ChAT^–lCeA^PKCδ^ pathway and eLPB^ChAT^–ovBNST^PKCδ^ pathway, respectively.

In summary (Fig. 4), our findings for the first time identified eLPB^ChAT^– lCeA^PKCδ^ pathway and eLPB^ChAT^–ovBNST^PKCδ^ pathway, and revealed the involvement of synaptic elements of presynaptic Ach release and postsynaptic nAChRs in the positive innervation of these cholinergic pathways. Importantly, METH withdrawal anxiety and METH-primed reinstatement of CPP recruit eLPB^ChAT^–lCeA^PKCδ^ and eLPB^ChAT^–ovBNST^PKCδ^ pathway in METH-exposed male mice, respectively.

## Online content

Methods, extended data, supplementary information, acknowledgements, peer review information; details of author contributions and competing interests; and statements of data and code availability will be available online when it is accepted.

## References

1. Su, H., et al. Anxiety level and correlates in methamphetamine-dependent patients during acute withdrawal. Medicine 96, e6434 (2017).

2. Xu, X., et al. Specific Inhibition of Interpeduncular Nucleus GABAergic Neurons Alleviates Anxiety-Like Behaviors in Male Mice after Prolonged Abstinence from Methamphetamine. The Journal of neuroscience : the official journal of the Society for Neuroscience 43, 803–811 (2023).

3. Campillo, R. My Experience and Recovery from Meth Addiction. Missouri medicine 119, 500 (2022).

4. He, T., et al. A novel cholinergic projection from the lateral parabrachial nucleus and its role in methamphetamine-primed conditioned place preference. Brain Commun 4, fcac219 (2022).

5. Coronel-Oliveros, C., Cofré, R. & Orio, P. Cholinergic neuromodulation of inhibitory interneurons facilitates functional integration in whole-brain models. PLOS Computational Biology 17, e1008737 (2021).

6. Picciotto, M.R., Higley, M.J. & Mineur, Y.S. Acetylcholine as a neuromodulator: cholinergic signaling shapes nervous system function and behavior. Neuron 76, 116–129 (2012).

7. Li, X., et al. Generation of a whole-brain atlas for the cholinergic system and mesoscopic projectome analysis of basal forebrain cholinergic neurons. Proceedings of the National Academy of Sciences 115, 415–420 (2018).

8. Seiglie, M.P., Lepeak, L., Miracle, S., Cottone, P. & Sabino, V. Stimulation of lateral parabrachial (LPB) to central amygdala (CeA) pituitary adenylate cyclase-activating polypeptide (PACAP) neurons induces anxiety-like behavior and mechanical allodynia. Pharmacology Biochemistry and Behavior 230, 173605 (2023).

9. Liu, S., et al. Divergent brainstem opioidergic pathways that coordinate breathing with pain and emotions. Neuron 110, 857–873.e859 (2022).

10. Boucher, M.N., Aktar, M., Braas, K.M., May, V. & Hammack, S.E. Activation of Lateral Parabrachial Nucleus (LPBn) PACAP-Expressing Projection Neurons to the Bed Nucleus of the Stria Terminalis (BNST) Enhances Anxiety-like Behavior. Journal of molecular neuroscience : MN 72, 451–458 (2022).

11. Jaramillo, A.A., Williford, K.M., Marshall, C., Winder, D.G. & Centanni, S.W. BNST transient activity associates with approach behavior in a stressful environment and is modulated by the parabrachial nucleus. Neurobiology of stress 13, 100247 (2020).

12. Venniro, M., et al. Abstinence-dependent dissociable central amygdala microcircuits control drug craving. Proceedings of the National Academy of Sciences of the United States of America 117, 8126–8134 (2020).

13. Domi, E., et al. Activation of GABAB receptors in central amygdala attenuates activity of PKCδ + neurons and suppresses punishment-resistant alcohol self-administration in rats. Neuropsychopharmacology : official publication of the American College of Neuropsychopharmacology 48, 1386–1395 (2023).

