## Supplemental Figures for "Distinct eLPB^ChAT^ projections for methamphetamine anxiety and relapse"

#### Extended Data Fig. 1

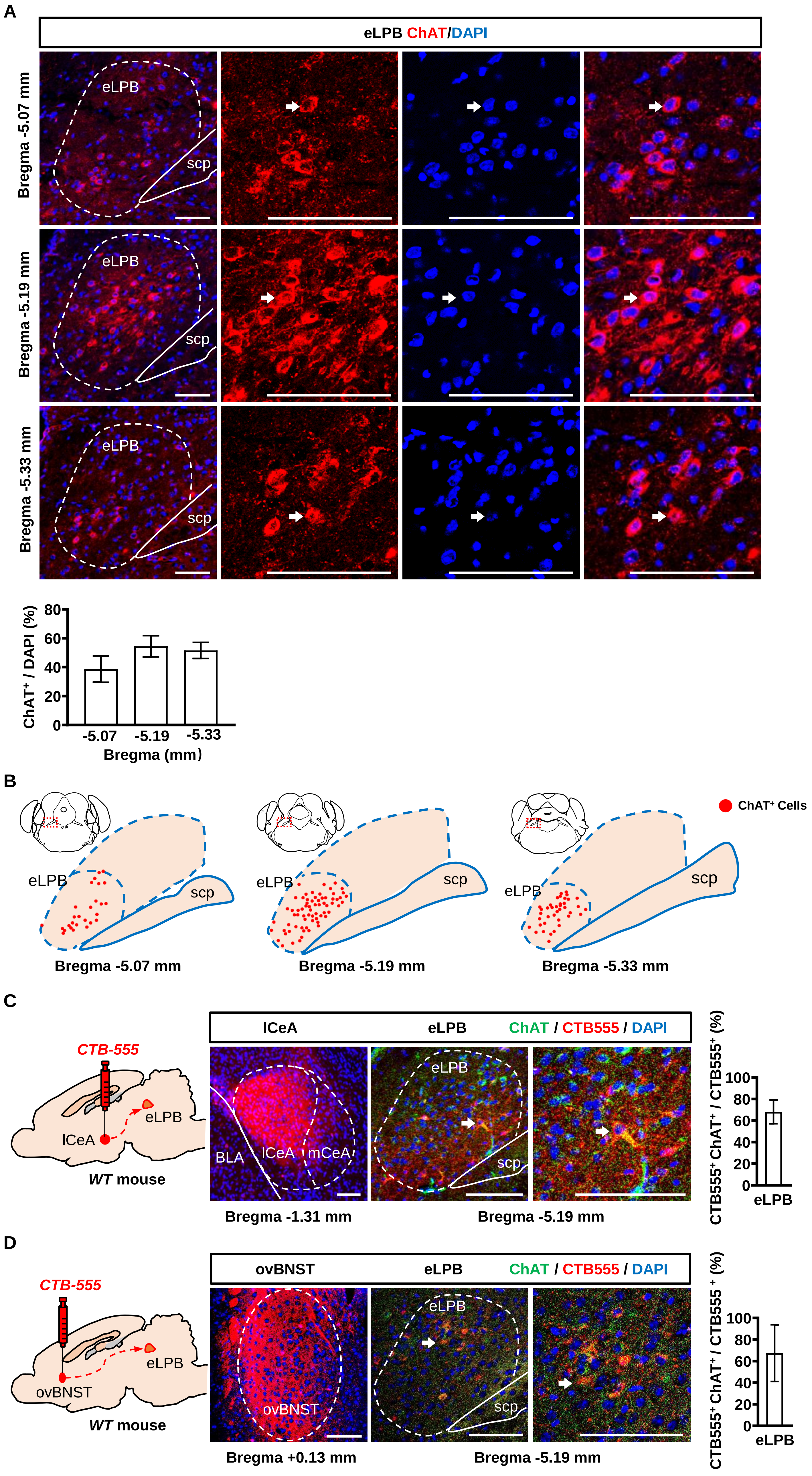

### Extended Data Fig. 2

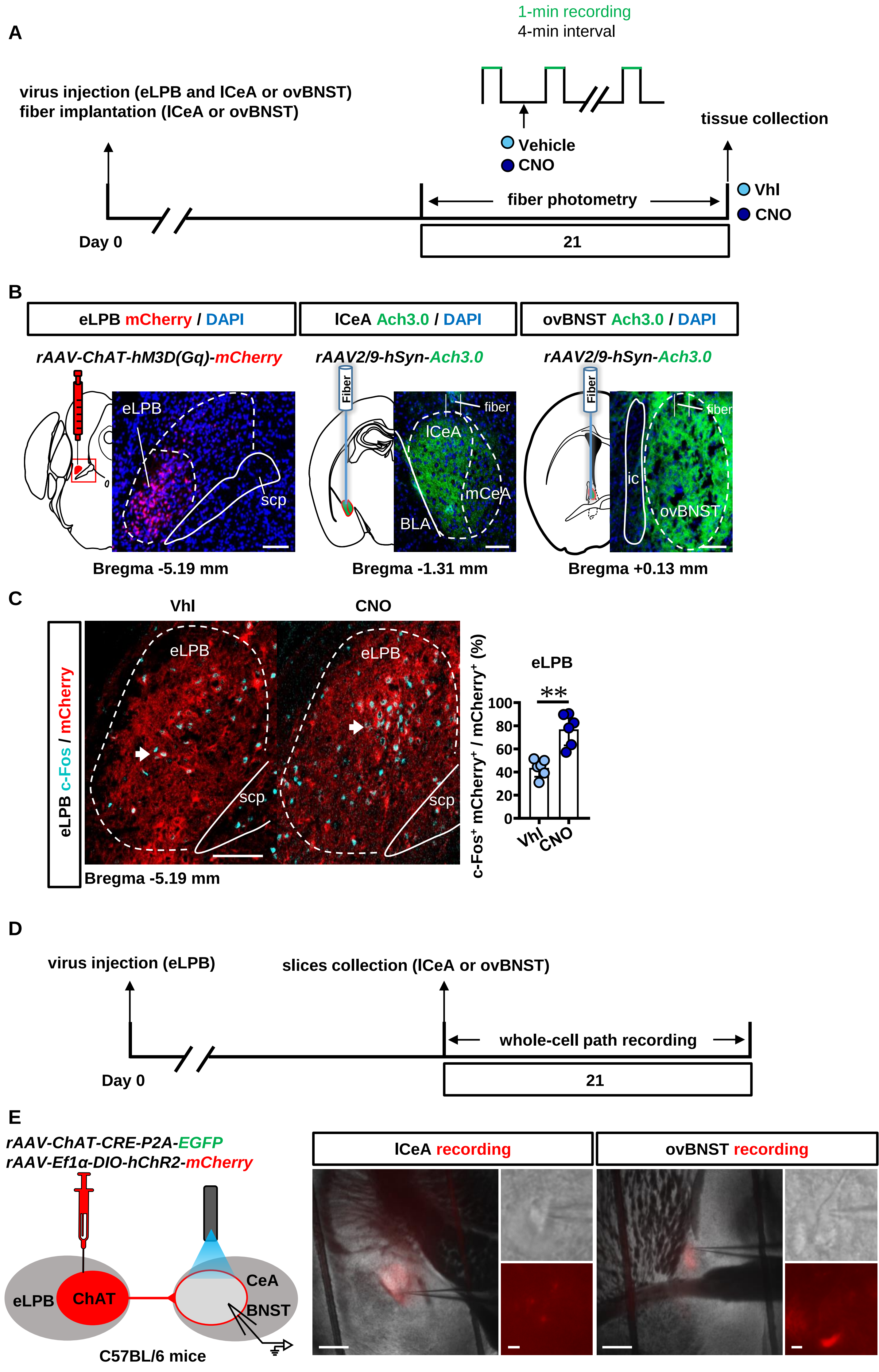

### Extended Data Fig. 3

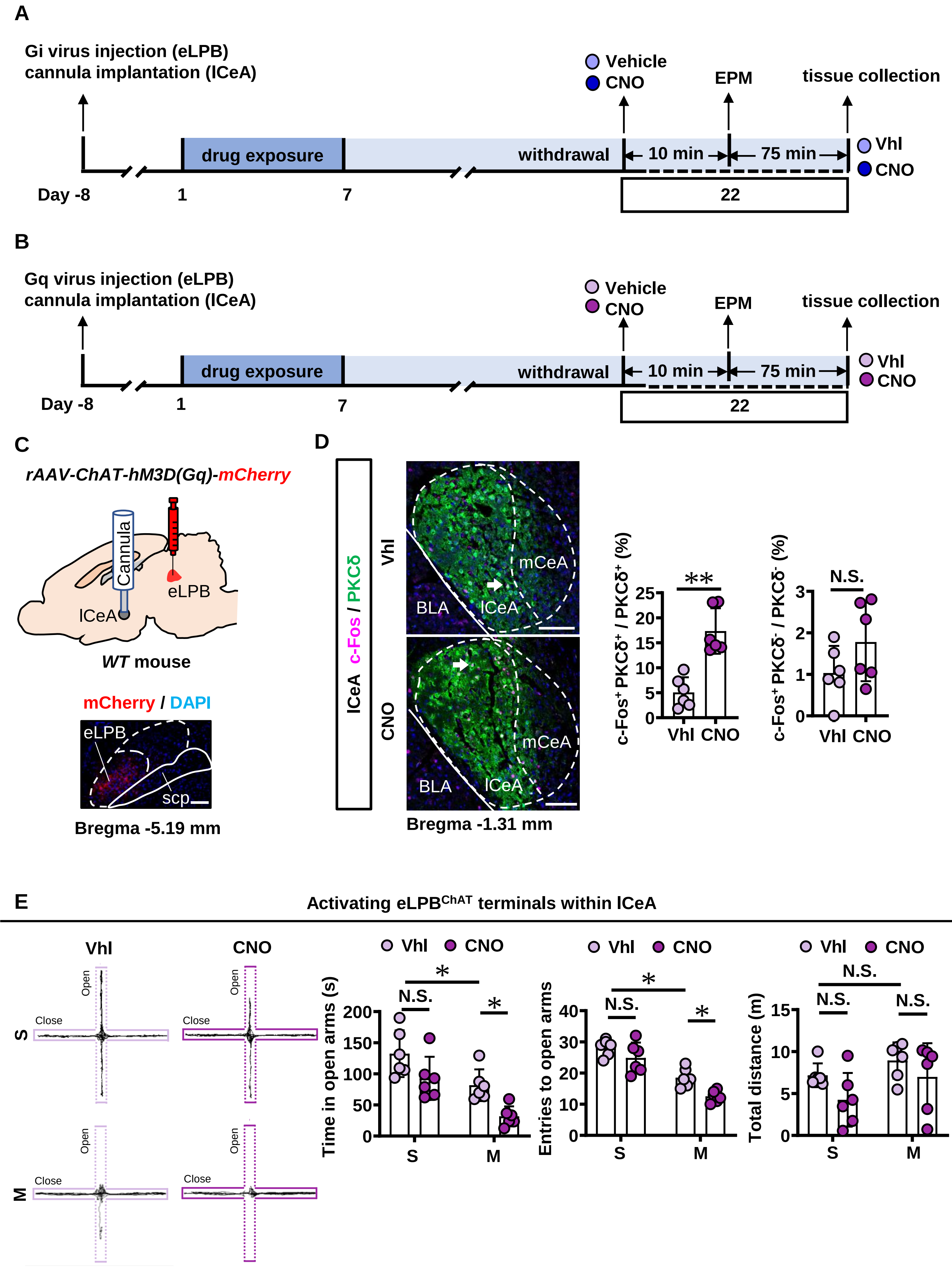

#### Extended Data Fig. 4

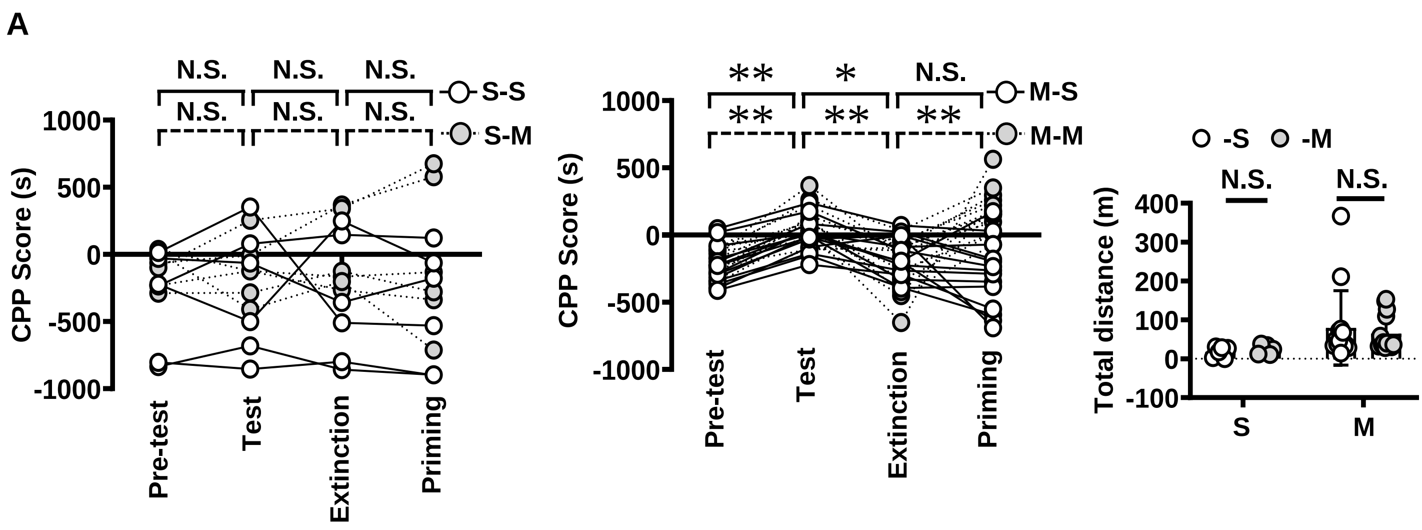

**Extended Data Fig. 5**

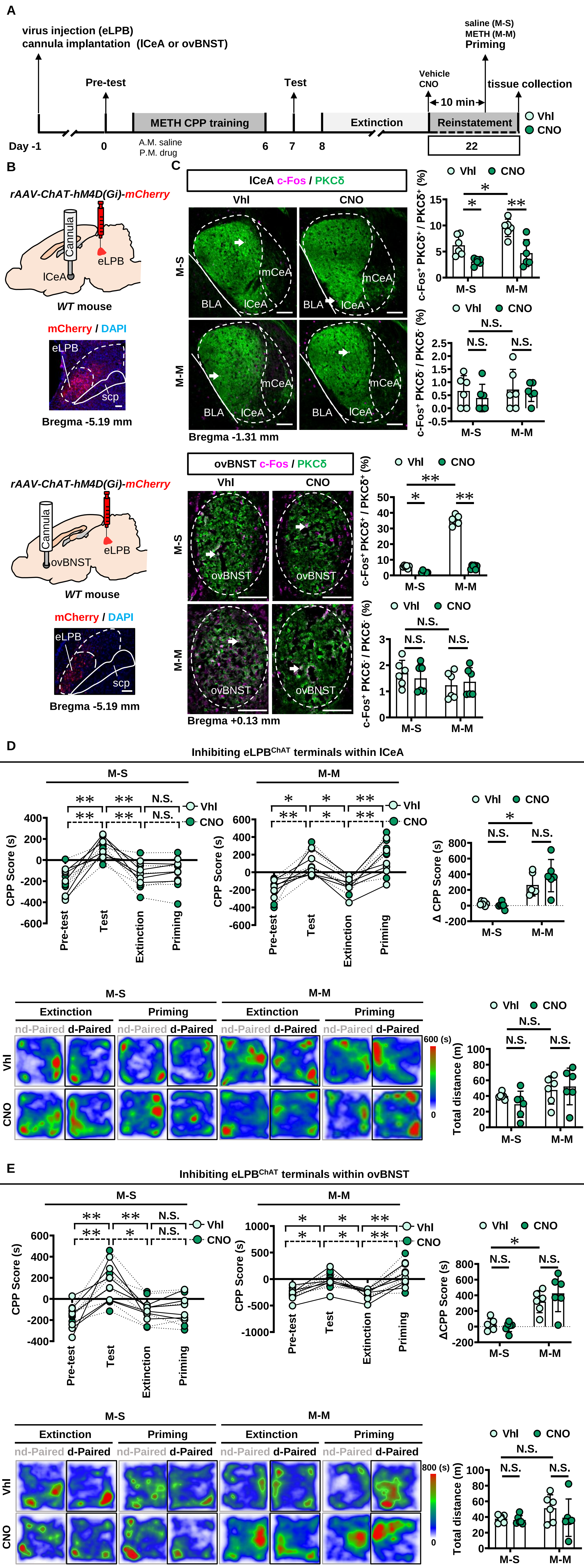

### Extended Data Fig. 6

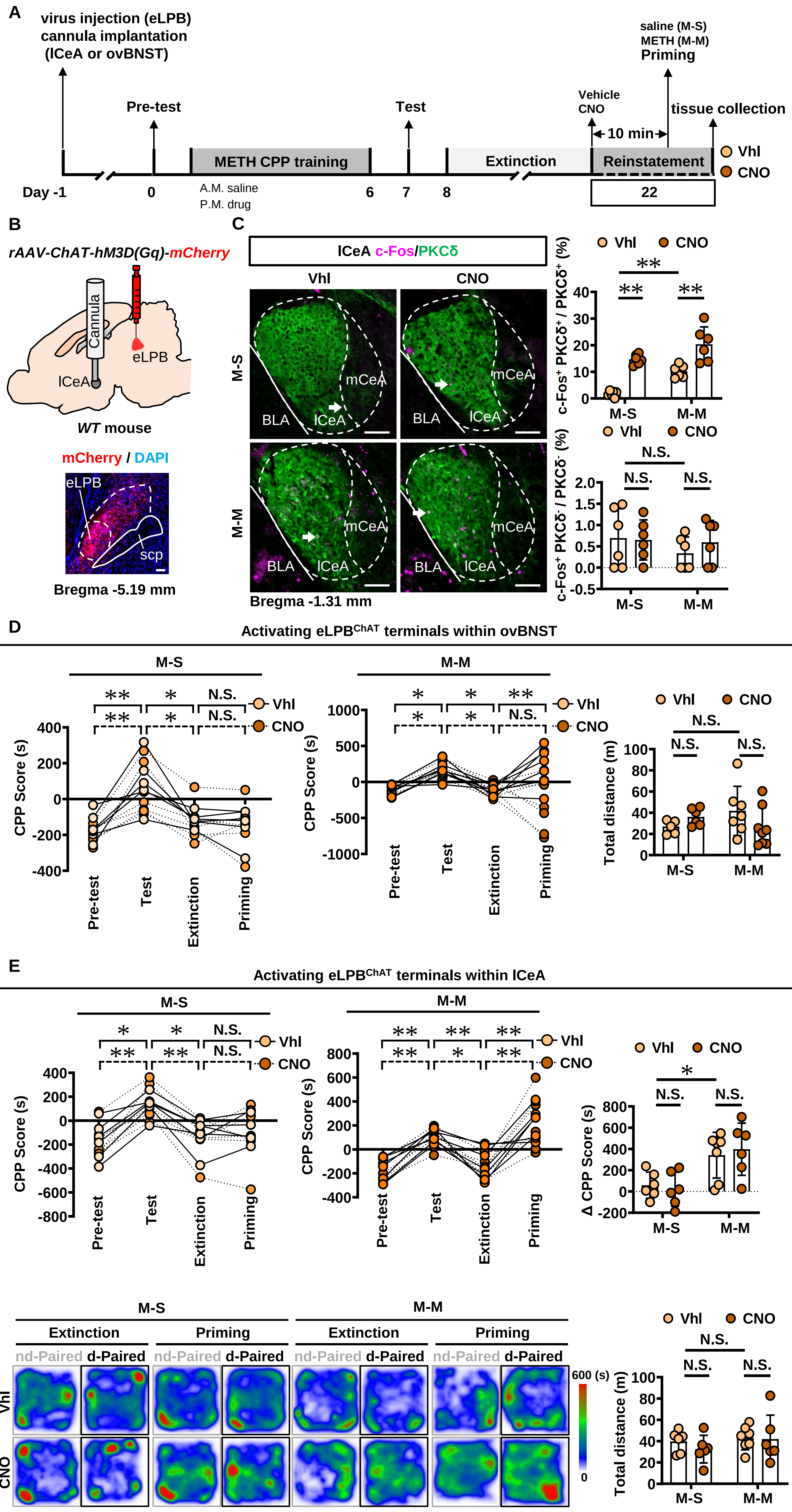
